# A Deep Graph Neural Network Architecture for Modelling Spatio-temporal Dynamics in resting-state functional MRI Data

**DOI:** 10.1101/2020.11.08.370288

**Authors:** Tiago Azevedo, Alexander Campbell, Rafael Romero-Garcia, Luca Passamonti, Richard A.I. Bethlehem, Pietro Liò, Nicola Toschi

## Abstract

Resting-state functional magnetic resonance imaging (rs-fMRI) has been successfully employed to understand the organisation of the human brain. For rs-fMRI analysis, the brain is typically parcellated into regions of interest (ROIs) and modelled as a graph where each ROI is a node and pairwise correlation between ROI blood-oxygen-level-dependent (BOLD) time series are edges. Recently, graph neural networks (GNNs) have seen a surge in popularity due to their successes in modelling unstructured relational data. The latest developments with GNNs, however, have not yet been fully exploited for the analysis of rs-fMRI data, particularly with regards to its spatio-temporal dynamics. Herein we present a novel deep neural network architecture, combining both GNNs and temporal convolutional networks (TCNs), which is able to learn from the spatial and temporal components of rs-fMRI data in an end-to-end fashion. In particular, this corresponds to intra-feature learning (i.e., learning temporal dynamics with TCNs) as well as inter-feature learning (i.e., leveraging spatial interactions between ROIs with GNNs). We evaluate our model with an ablation study using 35,159 samples from the UK Biobank rs-fMRI database. We also demonstrate explainability features of our architecture which map to realistic neurobiological insights. We hope our model could lay the groundwork for future deep learning architectures focused on leveraging the inherently and inextricably spatio-temporal nature of rs-fMRI data.

## 1. Introduction

Resting-state functional magnetic resonance imaging (rs-fMRI) is one of the most commonly used noninvasive imaging techniques employed to gain insight into human brain function. The use of rs-fMRI data has proven extremely useful as an investigative tool in neuroscience and, to some extent, as a biomarker of brain disease diagnosis and progression [1]. Typical use of rs-fMRI data involves using graph-theoretical measures (such as centrality measures and community structure) to summarise high-dimensional, whole-brain data for use in downstream tasks. As part of this process, it is common practice to reduce the dimensionality of the data in one of three main ways: (1) by collapsing the temporal dimension (e.g., into brain region connectivity matrices based on similarity metrics between time series), (2) by reducing the spatial dimension (e.g., in global signal regression for physiological noise modelling [2]), and (3) by employing approaches that collapse both the temporal and spatial dimensions (e.g., in independent component analyses [3]). This feature engineering step is performed mostly due to the considerable volume of data in a typical rs-fMRI dataset and its relatively low signal-to-noise ratio [4]. Although computationally beneficial, such dimensionality reduction steps inevitably involve disregarding large amounts of information which can potentially be useful depending on the analysis task. For instance, collapsing the temporal dimension of rs-fMRI data reduces the brain to a static volume where the interactions between different brain regions are fixed over time. This stands in contrast to a growing body of research showing that the functional connectivity of the brain is dynamic and constantly changing over time [5, 6]. As another example, association measures most commonly used are still based on linear models, while it is well known that neuromonitoring data and brain signal in particular interact nonlinearly [7, 8].

To overcome such limitations, a different approach to the analysis of rs-fMRI data would be to devise a model that is able to combine both feature engineering and the learning of a low-dimensional representation of the brain. In order to do this, such a model would need to be able to accommodate both the spatial as well as the temporal complexities of rs-fMRI data. To date, deep learning architectures have had great success at leveraging specific inductive biases from complex high-dimensional data. Convolutional neural networks (CNNs), for instance, are extremely effective at extracting shared spatial features such as corners and edges from grid-like data (e.g., 2D and 3D images). These features can then be combined into more complex concepts deeper within the network architecture [9]. Recurrent neural networks (RNNs), on the other hand, are able to learn features from data that are temporally organised as a sequence of steps [10]. In contrast to both CNNs and RNNs, graph neural networks (GNNs) can learn from data that does not have a rigid structure like a grid or a sequence, and can be depicted in the form of unordered entities and relations such as graphs. The formulation of GNN models that deal with complex data structures has seen fast developments in the past years [11, 12] and are therefore strong candidates for the analysis of rs-fMRI data.

Previous work has attempted to leverage deep learning architectures in order to model rs-fMRI data. In particular, GNNs have been used to classify binary sex [13], and CNNs have been successfully employed in the diagnosis of cognitive impairment [14]. Other pioneering studies have devised *ad-hoc* deep learning models for fMRI data such as the classification of brain disorders using Siamese-inspired neural networks [15], but the spatial information was not represented using GNNs. Learning from spatial and temporal components of data using deep learning can also be seen in various non-biological domains [16, 17, 18]; such approaches, however, usually rely on a single spatio-temporal convolutional block that creates low-dimensional embeddings at each timestep, instead of performing a prediction task for the entire graph at once. To improve these drawbacks, we formulate a novel deep neural network architecture that exploits the advantages of GNNs and CNNs in order to effectively model the linear and non-linear temporal and spatial components of rs-fMRI data. We engineered our architecture to specifically retain edge weights and contain elements of explainability [19, 20], in order to provide advantages when a neuroscientific explanation of the inner model workings is desirable. Our proposed model uses GNNs to account for spatial inter-relationships between brain regions, and temporal convolutional networks (TCNs) to capture the intra-temporal dynamics of blood-oxygenated-level dependent (BOLD) time series. By incorporating GNNs and CNNs in the same end-to-end architecture we essentially combine intra- and inter-feature learning. In particular, GNNs can tackle a limitation of some graph representations of rs-fMRI data, in which association measures between different regions of interest (ROIs) of the brain are based on linear models; instead, GNNs can capture higher-order interactions between ROIs. A very preliminary version of this work with a 30-fold smaller dataset was recently presented as a conference contribution [21]. However, further work was needed regarding a larger dataset, wider choices of the graph threshold hyperparameter, and analysis on inclusion of edge weights.

We test our architecture on the publicly available UK Biobank dataset, which at the time of writing provides rs-fMRI scans from more than 30,000 distinct people. This dataset offers a unique opportunity to formulate novel architectures, while supporting the need of large datasets for reproducible findings with minimal statistical errors [22]. We also conducted an ablation analysis on a proof-of-concept binary sex prediction task to better evaluate the different contributions of each component of our model. We release all the code and artifacts used to develop this work in a public repository for easier adoption by the community (see “Data and Code Availability” section).

## 2. Methods

### 2.1. Problem Definition

To represent rs-fMRI data as an undirected weighted graph, the brain is spatially parcellated into *N* regions of interest (ROIs) representing graph nodes indexed by the set 𝒱= {1, …, *N*}. Let ***x***_*i*_ ∈ℝ^*T*^ represent the features of node *i* corresponding to the BOLD timeseries of length *T*. The connections between each ROI are represented by an edge set *ε* ⊂ 𝒁 × 𝒱composed of = *E* unordered pairs (*i, j*), where for every edge *k* connecting two nodes (*i, j*) ∈ *ε* the connection strength is defined as ***e***_*k*_ ∈ ℝ. Let the tuple 𝒢 = (𝒱, *ε*) denote the resulting graph. Given the graph structure 𝒢, let ***X*** ∈ ℝ^*N* ×*T*^, ***E*** ∈ ℝ^*E*×1^, and ***A*** ∈ ℝ^*N* ×*N*^ denote the nodes features, edge features and adjacency matrix, respectively.

### 2.2. Temporal Convolutional Networks

In order to learn a representation of the temporal dynamics contained in rs-fMRI time series, we use temporal convolutional networks (TCNs) [23]. These are a simplification over the original *WaveNet* architecture used for audio synthesis [24], which has been seen to provide significantly better results for sequence modelling in comparison to more traditional RNN architectures (e.g., LSTMs) across a range of tasks and datasets. In particular, Bai et al. [23] posit that convolutional networks should be seen as the natural starting point for sequence modelling tasks, which makes them ideal for extracting information from rs-fMRI time series.

TCNs are based on dilated causal convolutions [25], which are special 1D filters where the size of the receptive field exponentially increases over the temporal dimension of the data as the depth of the network increases. The padding of the convolution is ‘causal’ in the sense that an output at a specific time step is convolved only with elements from earlier time steps from the previous layers, thus preserving temporal order. More formally, given a single ROI timeseries ***x***_*i*_ ∈ ℝ^*T*^ and a filter ***f*** ∈ ℝ^*K*^, the dilated causal convolution operation of ***x*** with ***f*** at time *t* is represented as

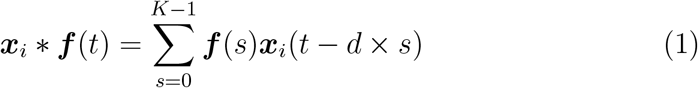

where *d* is the dilation factor which controls the number of time steps successively skipped. In contrast to the original TCN architecture [23], we use batch normalisation instead of weight normalisation because it empirically provided a more stable training procedure in terms of loss evolution.

### 2.3. Graph Network Block

Battaglia et al. [26] formalise a graph network (GN) framework through the definition of functions that work on graph-structured representations. The main unit of computation in the GN framework is called the *GN block*, which contains three update functions and three aggregation functions working on the edge, node, and global level.

The first operation of this GN block, which can be broadly defined as the *edge model*, concerns the update function *ϕ*^*e*^, which computes updated edge attributes for each edge *k* based on the original edge’s attributes ***e***_*k*_ and the features of the connected nodes *i* and *j*:

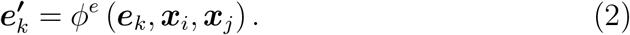

Note that for rs-fMRI graph representations, each edge originally contains a single value (i.e., ***e***_*k*_ ∈ ℝ), but after this operation *ϕ*^*e*^, the resulting dimensionality can be different: 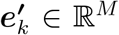, where *M* >= 1. Then, in what can be broadly defined as the *node model*, the block computes updated node features. Firstly, for each node *i*, it aggregates the edge features per node:

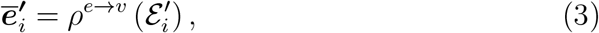

where 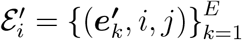 is the set of edges starting in node *i*, with node *j* connected with node *i* through edge *k*. Importantly, *ρ*^*e*→*v*^ needs to be invariant to edge permutations to account for the unordered structure of the data. Averaging and summation are examples of such operations invariant to edge permutations.

Finally, the updated node features are computed using another aggregation function at the node level, for each node *i*:

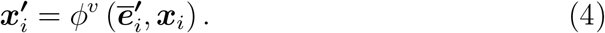

Although the rs-fMRI graph representation contains undirected edges, the GN block requires directed edges. To overcome this issue, every time there is a connection between any two nodes *i* and *j*, we assume the existence of two edges (***e***_*k*_, *i, j*) and (***e***_*k*_, *j, i*), one for each direction.

### 2.4. Graph Pooling

After the neural network processes the input as described in the previous sections, each node in the graph will contain a node-wise representation (i.e., a feature vector) as a result. For the prediction task described in this paper, where a graph-level (as opposed to node-level) prediction is required, these representations need to be pooled (i.e., collated) to be used for a final downstream prediction task.

To this end, it is common practice to employ a global average pooling mechanism, in which the nodes features are averaged across the graph, thus creating a final, low-dimension embedding representation of the graph itself.

However, assuming that distinct nodes (i.e., brain regions in this work) have different levels of importance for the downstream prediction task [27, 28], we assumed that a hierarchical (as opposed to flat) pooling mechanism would create richer embeddings. To this end, we employ the differentiable pooling operator introduced by Ying et al. [29], commonly called DiffPool, which learns how to sequentially collapse nodes in smaller clusters until only a single node exists with the final embedding.

When describing a Graph Network (GN) block, a sparse representation of nodes and edges is used to describe the operations that a GN block can have; however, DiffPool works on dense representations of a graph. In other words, a graph 𝒢 is represented by a dense adjacency matrix ***A*** ∈ ℝ^*N* ×*N*^ and a feature matrix ***X*** ∈ ℝ^*N* ×*F*^, where *N* is the number of nodes and *F* the number of features in each node.

The DiffPool operator, at layer *l*, thus receives both an adjacency matrix and a node embedding matrix, and computes updated versions of both:

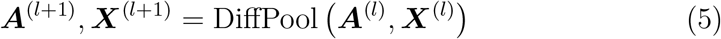

To achieve this, the DiffPool operator uses a graph neural network (GNN) architecture. Specifically, the same GNN architecture is duplicated to compute two distinct representations: a new embedding 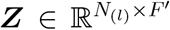 and an assignment matrix 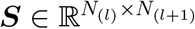:

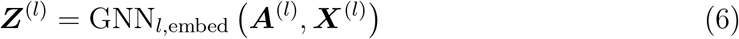

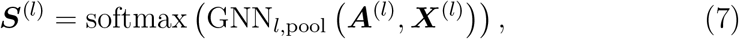

where *N*_(*l*)_ is the number of nodes in layer *l, N*_(*l*+1)_ the new number of nodes, each corresponding to a cluster (*N*_(*l*+1)_ < *N*_(*l*)_), and *F* ′ the number of features per node, which can be different from the original size *F* from the matrix ***X***.

The operator ends with the creation of the new node embedding matrix and adjacency matrix, to be inputted to the next layer:

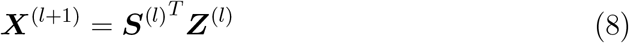

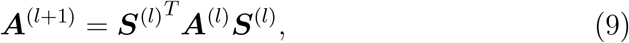

where 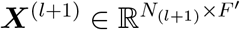 and 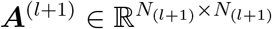.

## 3. Experiments Overview

### 3.1. Dataset UK Biobank

Subject-level structural T1 and T2-FLAIR data as well as ICA-FIX [30] denoised rs-fMRI data was obtained from UK BioBank (application 20904) [31]^1^. All data were acquired on a standard Siemens Skyra 3T scanner running VD13A SP4, with a standard Siemens 32-channel RF receive head coil. The structural data was further preprocessed with Freesurfer (v6.0)^2^ using the T2-FLAIR weighted image to improve pial surface reconstruction, similarly to Glasser et al. [32]’s pipeline. Reconstruction included bias field correction, registration to stereotaxic space, intensity normalisation, skull-stripping, and white matter segmentation. When no T2-FLAIR data was available, Freesurfer reconstruction was done using the T1 weighted image only. Following surface reconstruction, the Desikan-Killiany atlas [33] was aligned to each individual structural image and and ROIs extracted. The same atlas was aligned to the functional denoised rs-fMRI data (490 volumes TR/TE = 735/39.00 ms, multiband factor 8, voxel size: 2.4×2.4×2.4, FA=52 deg, FOV 210×210 mm) using the warping parameters from the structural to functional alignment obtained using FSL’s linear registration (FLIRT), and mean BOLD time series (490 timepoints per scan) were extracted for each ROI. The time series were then scaled subject-wise using the median and interquartile range according to the *RobustScaler* implementation from the *scikit-learn* [34] python package. Edge weights were defined as full correlations calculated with the Ledoit Wolf covariate estimator using the *nilearn* python package^3^. Figure 1 shows an example scaled time series and the resulting example graph from a single subject. The total number of subjects used from the UK Biobank was 35,159, in which 18,649 were females and 16,510 were males (18, 649*/*16, 510 ≈ 1.13). The median age was 64 with a minimum age of 44 and a maximum of 81.

**Figure 1:**
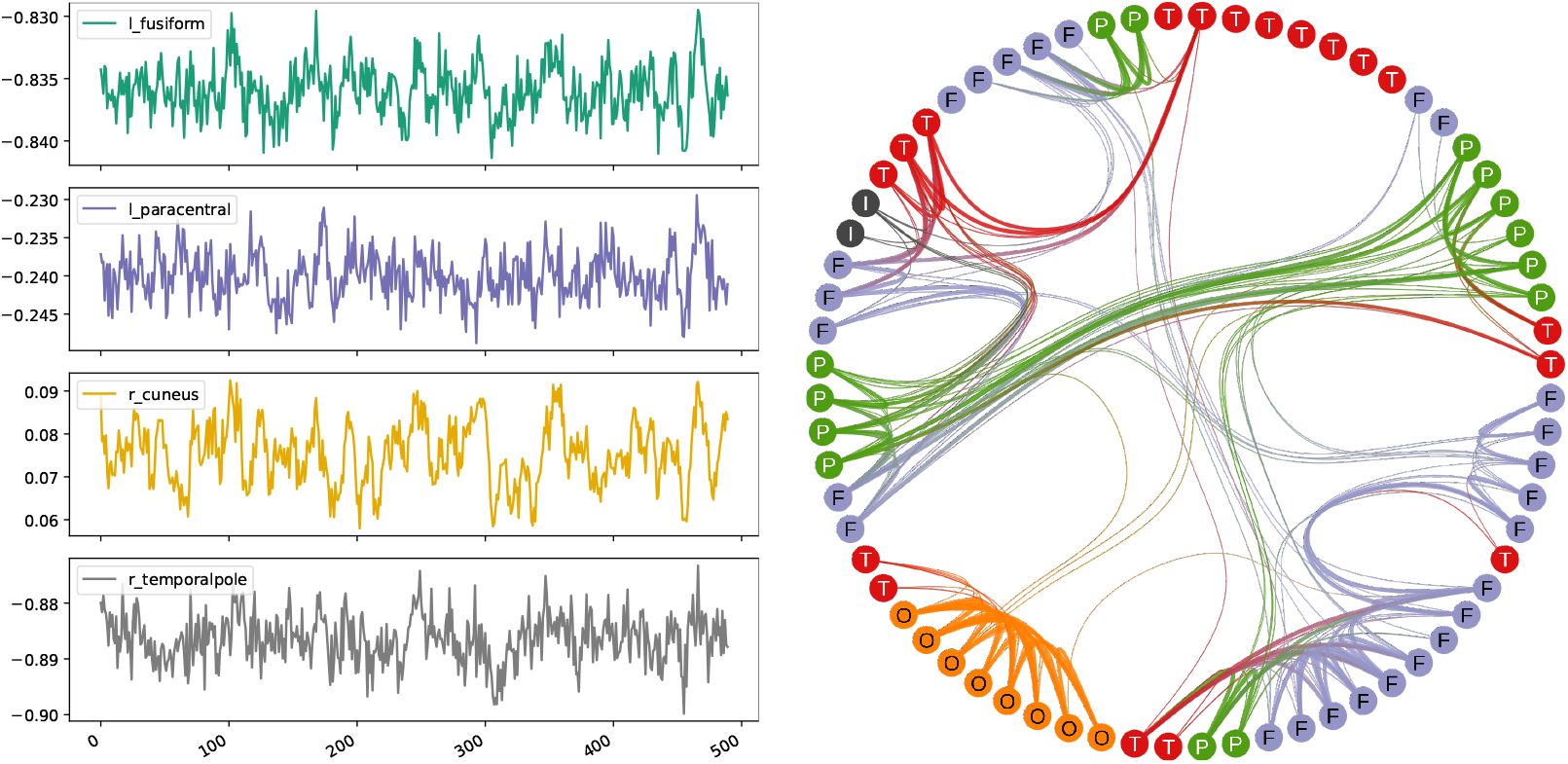
Left: Mean BOLD time series of four brain regions from the same subject, after scaling. Right: Graph representation of one subject’s data, at 10% threshold as described in Section 3.2. Thicker edges represent a stronger correlation between nodes, in this case with values between approximately 0.54 and 0.87. Each node is labelled and coloured according to its brain region (i.e., T/F/O/P/I correspond to <u>T</u>emporal, <u>F</u>rontal, <u>O</u>ccipital, <u>P</u>arietal, and Insula).

### 3.2 Model Implementation

The neural network architecture depicted in Figure 2 was implemented using Pytorch [35], and Pytorch Geometric [36] for the specific graph neural network components. The edge feature matrix ***E*** ∈ ℝ^*E*×1^ defined in Section 2.1 was implemented as two sparse matrices: a sparse representation of the adjacency matrix **E**_*i*_ ∈ ℝ^2×*E*^, and a sparse representation of the edge features **E**_*a*_ ∈ ℝ^*E*×1^ (i.e., there was only one feature per edge corresponding to the correlation value). The number of nodes *N* was 68 (corresponding to each brain region from the Desikan-Killiany atlas), the number of node features *F* was the number of timepoints (i.e., 490), and *E* is the number of edges in the graph. The number of edges depends on the threshold percentage used to retain only the strongest correlations. Given the non-conclusive evidence on the optimal threshold percentage in the vast majority of functional connectivity literature [37], in this work this threshold was included in the hyperparameters to be optimised.

**Figure 2:**
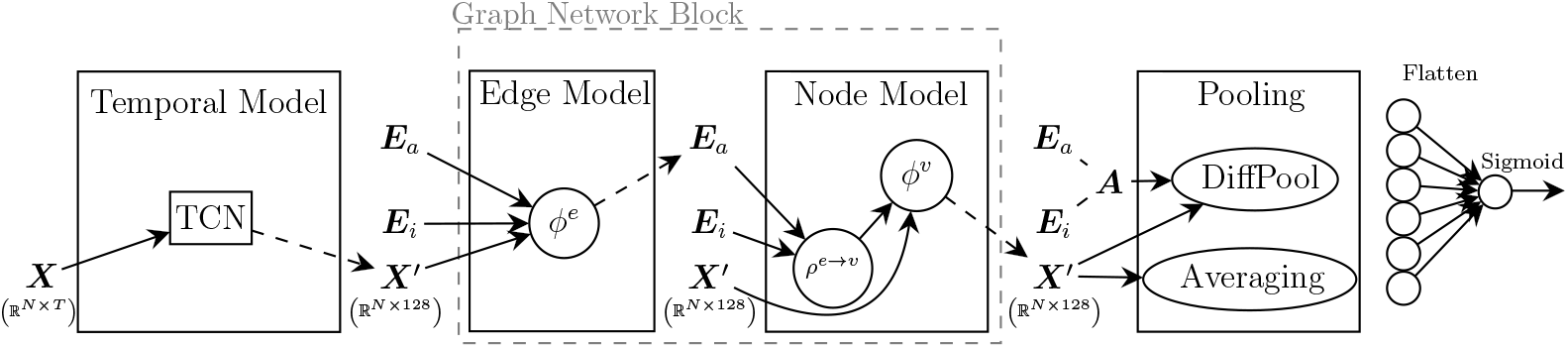
Three main working blocks of the spatio-temporal model. The temporal model creates an initial representation from original node features ***X*** (i.e., temporal dynamics). It is followed by transformations in the Graph Network Block which leverages the structure of data represented in edge features ***E***_*a*_ and sparse connectivity ***E***_*i*_. Finally, a Pooling mechanism (either DiffPool or global averaging) creates a final graph representation which is flatten for a final prediction task.

The full list of hyperparameters to be optimised and respective value range was the following:

- `dropout:` [0, 0.9] (uniform distribution)
- `threshold:` {5, 10, 20, 30, 40} (categorical)
- `learning rate:` [ln(1e−7), ln(1e−2)] (log uniform distribution)
- `weight decay:` [ln(1e−7), 0] (log uniform distribution)

To extract information from the rs-fMRI time series in each node, our model starts by employing a strided temporal convolutional network (TCN) architecture. Such architecture was implemented by using two blocks where each contained two layers of *1D convolutions, 1D batch normalisation, ReLU activation*, and *dropout*. Each block used a stride of 2, a kernel with size 7 (i.e., *K* = 7 in Equation 1), and contained a skip connection. While the first block used no dilation, the second one used a dilation of 2. The four *1D convolution* filters increased the numbers of output channels at each layer, specifically 8, 16, 32, and 64. After these two blocks (i.e., four layers), node features from all channels are flattened out and inputted to a linear transformation to reduce each node representation to a fixed embedding of size 128. These transformations thus reduce the original node feature matrix from size *N* × *T* to size *N* × 64 × 31 after the two blocks, and finally to size *N* × 128 corresponding to the final embedding.

Then, the Graph Network (GN) block is applied, in which the update functions are single-layered multi-layer perceptrons (MLPs), and aggregation function is an edge-wise averaging around each node. The original dimensions of ***X, E***_*i*_, and ***E***_*a*_ right before the GN block are kept after these transformations.

We employed two types of pooling mechanisms in our analysis, both of which reduce the node feature matrix from a size of *N* ×128 to 1 × 128: a global average pooling mechanism, and the hierarchical pooling mechanism (i.e., DiffPool). For DiffPool, which expects a dense graph representation, data is first transformed into a symmetric adjacency matrix ***A*** ∈ ℝ^*N* ×*N*^, which is a weighted matrix when considering edge features, and binary otherwise. As recommended by the original paper [29], we employed three layers of GraphSAGE [38] followed by a *1D batch normalisation*, with a final skip connection.

A summary conceptual architecture of the whole model is shown in Figure 2.

### 3.3. Training Procedure

In order to assess the validity of our model, we performed a proof-of-concept through the well-known binary sex prediction task [39, 40]. We used a 5-fold stratified cross validation procedure: the UK Biobank dataset was divided into training and test sets five times, in which each test set corresponds to 20% of the original size, and a sample would only belong to a test set once (i.e., all test sets are mutually exclusive). This division was done in a stratified fashion considering the sex label, bucketised age, and bucketised BMI measures (for each variable we created 8 equal-sized buckets based on sample quantiles). For each test set, the training set is further divided once to generate a single inner training and validation sets, using the same stratification strategy as for the training/test case.

The neural network was trained over 100 epochs with the Adam optimiser [41] and Binary Cross Entropy loss function. The training procedure was set to stop earlier if the validation loss did not reduce further after 33 consecutive epochs. A hyperparameter search was included in the inner training/validation sets, in which 25 random runs were launched exploring random values of dropout, edge threshold, learning rate, and weight decay (see Section 3.2 for values range). In each random run, the model with the smallest validation loss was saved, and the model with the smallest validation loss across the 25 runs was selected to be evaluated in the test set. This procedure is done separately for each test set, and metrics are averaged across the five test sets.

We used *Weights & Biases* [42] to log our training procedure and generate the random hyperparameters for all the 25 models in each inner sweep. These inner sweeps were run across two different servers, and each model took between 20 minutes and 11 hours to train depending on GPU type and early stopping. All these details are stored using *Weights & Biases*, and can be accessed through our public repository (see “Data and Code Availability”). Figure 3 shows the results for the inner sweep of one of the folds for illustrativeht2o2 purposes. While a certain amount of variability is visible, some trends are evident in this particular split: the best models (i.e., with lower validation loss) tend to be achieved with higher edge thresholds, higher learning rates, smaller weight decays, and smaller dropout rates. We highlight that different sweeps could result in very different trends.

**Figure 3:**
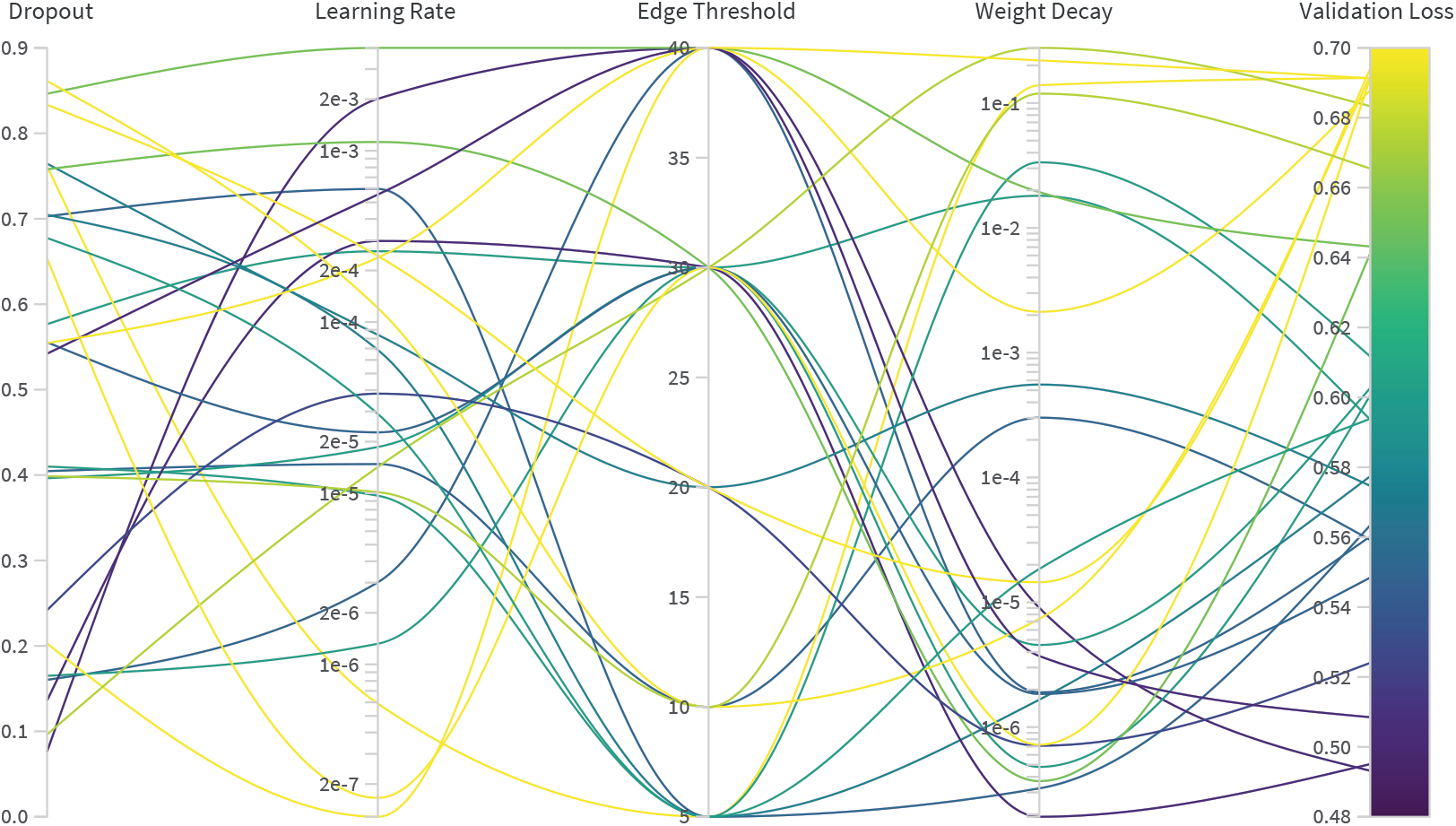
Values of hyperparameters corresponding to each validation loss achieved for one illustrative inner sweep of one fold. For each one of the 25 training runs (represented by each curved line), a set of random values is chosen for dropout, learning rate, edge threshold and weight decay, which ultimately will result in the model’s validation loss.

### 3.4. Evaluation

As shown in Figure 2, our model consists of (1) a TCN block that learns intra-temporal features from the mean BOLD time series of each ROI, followed by (2) a GN block which leverages the spatial inter-relationships between ROIs, and finally (3) a hierarchical pooling mechanism which leverages all the information in the input, from the temporal rs-fMRI dynamics, to the graph structure and the edge features of that graph.

In order to understand the inner workings of this combination, we conducted an ablation analysis to quantify the contributions of each component of our model for the specific prediction task. Firstly, we consider two cases where the GN block is not used, hence practically evaluating the importance of edge weights for this prediction task. In one case the graph structure is completely ignored (i.e., no GN block and global average pooling), and in another case a binary graph is used only for the final hierarchical pooling part (i.e., no GN block and DiffPool applied to a binary graph).

In order to investigate the influence of the different GN components, we consider not only the case where both *node model* and *edge model* are used in the GN Block, but also a case where only the *node model* is applied. For each one of these two cases, we consider both a global average pooling, as well as DiffPool with a weighted adjacency matrix.

## 4. Results

### 4.1. General Results

Table 1 shows the results of our ablation analysis across three different backbones - no graph block, only *node model*, and full graph network block each with two different aggregators (i.e., global average pooling and DiffPool). For notation purposes, we identify each one of these cases using “Backbone → Aggregator”, in which *Aggregator* can be “Average” or “DiffPool”, and *Backbone* can be “N” for only *node model*, “N + E” for both *node model* and *edge model* (i.e., full GN Block), and empty otherwise.

**Table 1:** Ablation analysis, with metrics averaged across the five test sets, with standard deviation in parenthesis. Aggregator on the right-hand side of the arrow, “N” corresponds to only *node model*, and “N + E” corresponds to full Graph Network block. **Params** stands for number of parameters.

| Model | AUC | Accuracy | Sensitivity | Specificity | Params |
| --- | --- | --- | --- | --- | --- |
| N + E → Average | 0.83 (0.013) | 0.76 (0.012) | 0.74 (0.035) | 0.77 (0.032) | 474,639 |
| N + E → DiffPool | 0.84 (0.009) | 0.76 (0.010) | 0.78 (0.042) | 0.74 (0.055) | 849,057 |
| N → Average | 0.80 (0.006) | 0.73 (0.007) | 0.71 (0.037) | 0.74 (0.030) | 441,486 |
| N → DiffPool | 0.84 (0.009) | 0.76 (0.008) | 0.77 (0.014) | 0.75 (0.016) | 815,904 |
| → DiffPool | 0.81 (0.021) | 0.73 (0.019) | 0.73 (0.035) | 0.73 (0.049) | 648,979 |
| → Average | 0.77 (0.004) | 0.70 (0.005) | 0.68 (0.004) | 0.73 (0.008) | 274,561 |

Using a GNN component achieves better results overall when compared with the “→ Average” case (i.e., no GNN), with clearer gains for AUC and accuracy metrics. Using DiffPool as an aggregator appears to deliver the best averaged metrics for AUC, accuracy, and sensitivity. The simplest GNN model in terms of number of parameters (i.e., “N → Average”) yields comparable results to other models, thus likely being the best compromise in terms of model complexity and performance power. Using the *edge model* did not bring significantly better results when compared to only using the *node model*, thus indicating that the information contained in the edge attributes is successfully leveraged by the *node model* alone for this particular prediction task.

The results presented so far consider an adjacency matrix threshold below 50% as a hyperparameter at training time, a common data reduction practice in the connectivity analysis field. We further analysed the results of using no threshold at all, and explored the type of activation function as a hyperparameter instead (i.e., *ReLU* or *tanh* activations). This choice was made explicitly since retaining 100% of the adjacency matrix elements results in a share of negative correlation elements, whose physiological significance is likely to be important in brain connectivity [43]. The results of this analysis are presented in Table 2.

**Table 2:** Results with no thresholded graphs, with metrics averaged across the five test sets, with standard deviation in parenthesis. Aggregator on the right-hand side of the arrow, “N” corresponds to only *node model*, and “N + E” corresponds to full Graph Network block. **Params** stands for number of parameters.

| Model | AUC | Accuracy | Sensitivity | Specificity | Params |
| --- | --- | --- | --- | --- | --- |
| N + E → Average | 0.79 (0.017) | 0.72 (0.013) | 0.71 (0.039) | 0.73 (0.027) | 474,639 |
| N + E → DiffPool | 0.79 (0.045) | 0.69 (0.077) | 0.55 (0.264) | 0.81 (0.098) | 849,057 |
| N → Average | 0.80 (0.032) | 0.72 (0.028) | 0.68 (0.077) | 0.76 (0.059) | 441,486 |
| N → DiffPool | 0.81 (0.009) | 0.74 (0.009) | 0.73 (0.060) | 0.74 (0.042) | 815,904 |

The performance was lower for all cases which did not implement a threshold. A possible explanation would be the excessive “noise” not allowing the dominating spatial structure of the graph to be successfully leveraged in a practical timeframe, possibly generating some overfitting. However, metrics are still comparable to the ones in Table 1, suggesting that these models are still able to extract some information from the data within only 100 epochs of training.

### 4.2. Explainability

Although deep neural networks are usually regarded as “black boxes”, in this paper we strived to inject some elements of explainability by inspecting selected learnt mechanisms after training. For instance, the weights of the TCN layers can be visually inspected. We visualised the first two layers of one of the trained N + E → DiffPool model trained on unthresholded matrices. For example, Figure 4 shows the weights learnt from the first TCN layer (each row corresponding to one of the 8 output channels of that layer), while Figure 5 depicts the same for the second TCN layer (each row corresponding to one of the 16 output channels and the columns corresponding to the 8 kernels of size 7 coming from the previous 8 channels).

**Figure 4:**
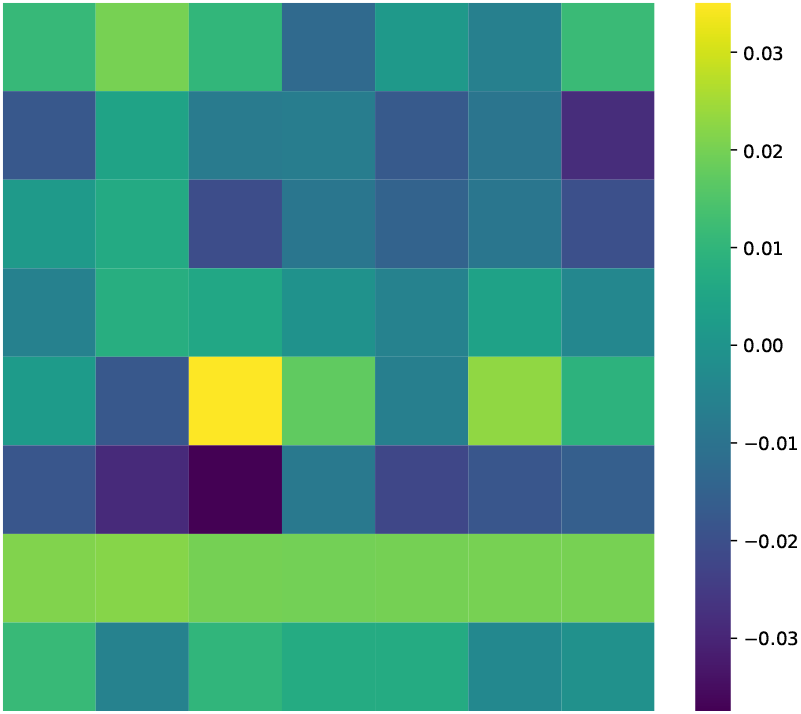
Weights of the kernels in the first TCN convolutional layer in a N + E → Diff-Pool model trained on unthresholded graphs. Rows correspond to the 8 output channels of this layer, and each column is a position in the kernel array of size 7

**Figure 5:**
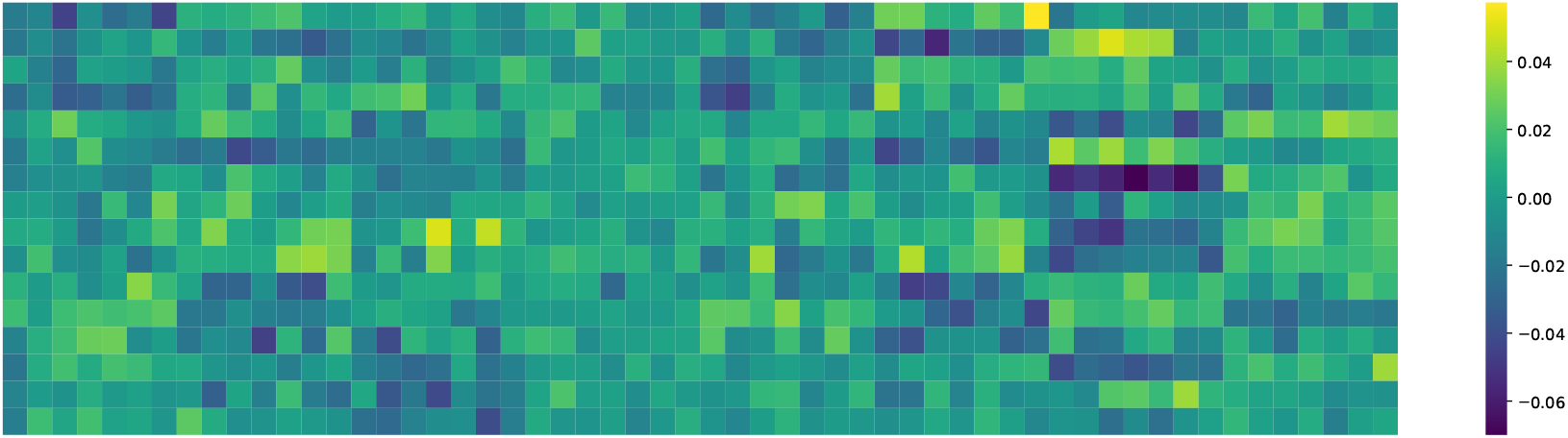
Weights of the kernels in the second TCN convolutional layer in a N + E → Diff-Pool model trained on unthresholded graphs. Rows correspond to the 16 output channels of this layer, and each column is a position in the 8 kernels of size 7 that come from the 8 input channels (56 columns in total).

In both figures, and with just a few exceptions, it can be seen that the output channels in the first two TCN convolutional layers are a nontrivial weighted multiplication of input channels. Given the qualitative variability observed in these learnt weights, we argue that these kernels are likely filtering and selecting different, non-mutually redundant patterns presented in the original time series. One possible counterexample is the kernel for the 7th output channel in the first TCN convolutional layer illustrated in Figure 4, which, in practice, is applying a simple low pass filter by smoothing the original time series from the input channel. We posit that quantitative analysis and comparison of the kernel weights across time series (which is out of the scope of the present paper), has the potential to yield interpretable information on which brain dynamics may contribute the most to the final prediction. It can also potentially highlight what frequency components (likely attached to different physiological significance [44]) are selected most often.

We further designed a strategy to inspect the hierarchical spatial pooling mechanism provided by the DiffPool architecture. To this end, we analysed the assignment matrices from the first Diffpool layer ***S***^(1)^ (see Equation 7), over all participants across all test sets. This is of particular interest because it corresponds to an aggregation of subsets of brain regions which our architecture has considered optimal while learning a particular prediction task. These aggregations can therefore be considered “optimal” for that task within this architecture, and provide hints to merge with neurophysiological knowledge. An assignment matrix corresponds to how the original nodes in the graph will be mapped into new nodes, thus, a direct way of summarising this effect across individuals is to count how many times two ROIs have ended up in the same cluster, regardless of cluster size and number. More formally, we create an association matrix ***S***′ ∈ ℝ^68×68^, where each element 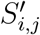 is the number of times brain regions *i* and *j* have been aggregated in the same DiffPool cluster. This means that the higher the value of 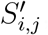, the more often information from brain regions *i* and *j* is pooled when learning to predict binary sex. It is important to point out that, given that matrix thresholding can potentially disconnect nodes from the rest of the graph and hence yield subject-wise graphs with variable numbers of connected nodes, this inspection strategy is only possible when working with unthresholded matrices (i.e., N → DiffPool and N + E → DiffPool models, see “Experiments Overview” section).

Figure 6 depicts the association matrix ***S***′ for the N + E → DiffPool model trained on unthresholded matrices, with dendrograms resulting from hierarchical clustering of the summary association matrix elements itself. For visualisation purposes in a more traditional brain connectivity style, we selected the four main clusters defined by the dendrograms for the N + E → DiffPool model and overlaid these clusters on a brain surface in Figure 7.

**Figure 6:**
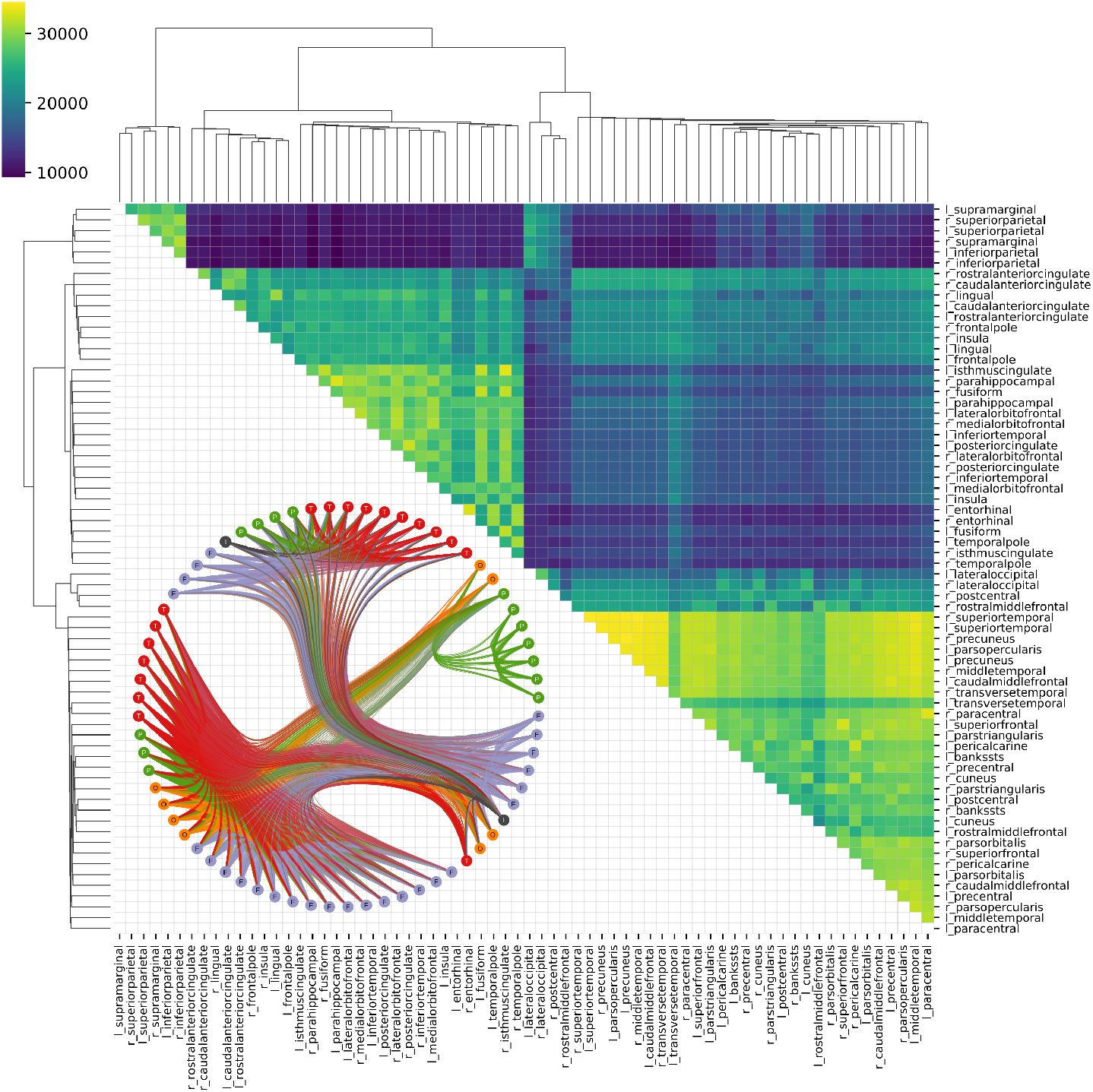
Upper-triangle of the association matrix **S**′ for N + E → DiffPool model generated when predicting binary sex on unthresholded matrices, with dendrograms from hierarchical clustering. Each element 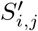 indicates how many times brain regions *i* and *j* are pooled together. On the lower left corner, a graph representation of the same association matrix **S**′, thresholded at 50% with nodes identified and coloured according to their general brain region (i.e., T/F/O/P/I correspond to <u>T</u>emporal, <u>F</u>rontal, <u>O</u>ccipital, <u>P</u>arietal, and Insula); thicker edges represent a higher 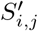 value, in this graph representation ranging from 20, 256 to 34, 565.

**Figure 7:**
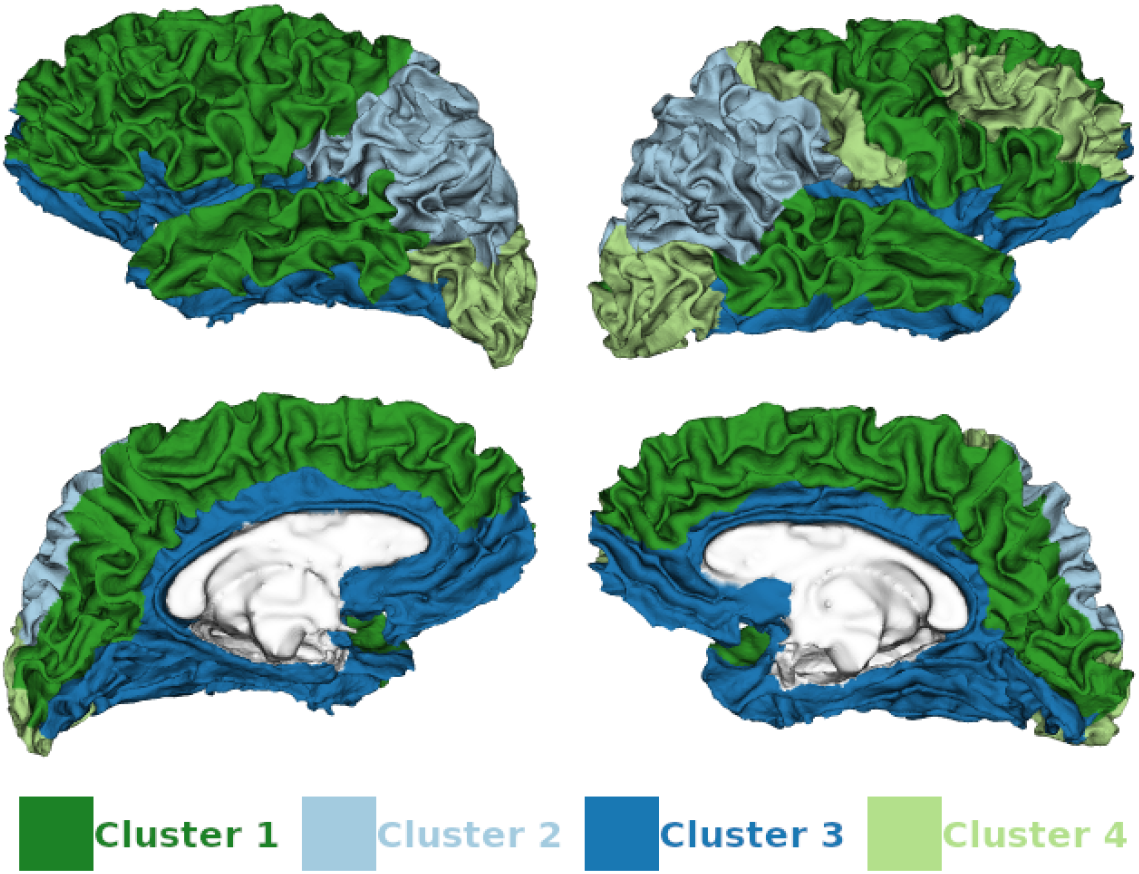
Four main brain clusters on association matrix ***S***′ generated from N + E → DiffPool model predicting binary sex on unthresholded matrices. Each colour corresponds to one cluster.

An advantage of an explainability constructing using the association matrix ***S***′ is the flexibility given to define the granularity used to select the clusters from the hierarchical clustering. When choosing big clusters (e.g., four like in Figure 7) one can illustrate the more general patterns, but by selecting smaller clusters (e.g. twelve clusters) one can reveal more local patterns in the data. We consider that these different levels of granularity are an advantage of using DiffPool to help explain the model, in practice revealing different scales of explainability. In Figure 8 we depict the brain clusters for the N + E → DiffPool model with the remaining different granularities (i.e., 8 and 12).

**Figure 8:**
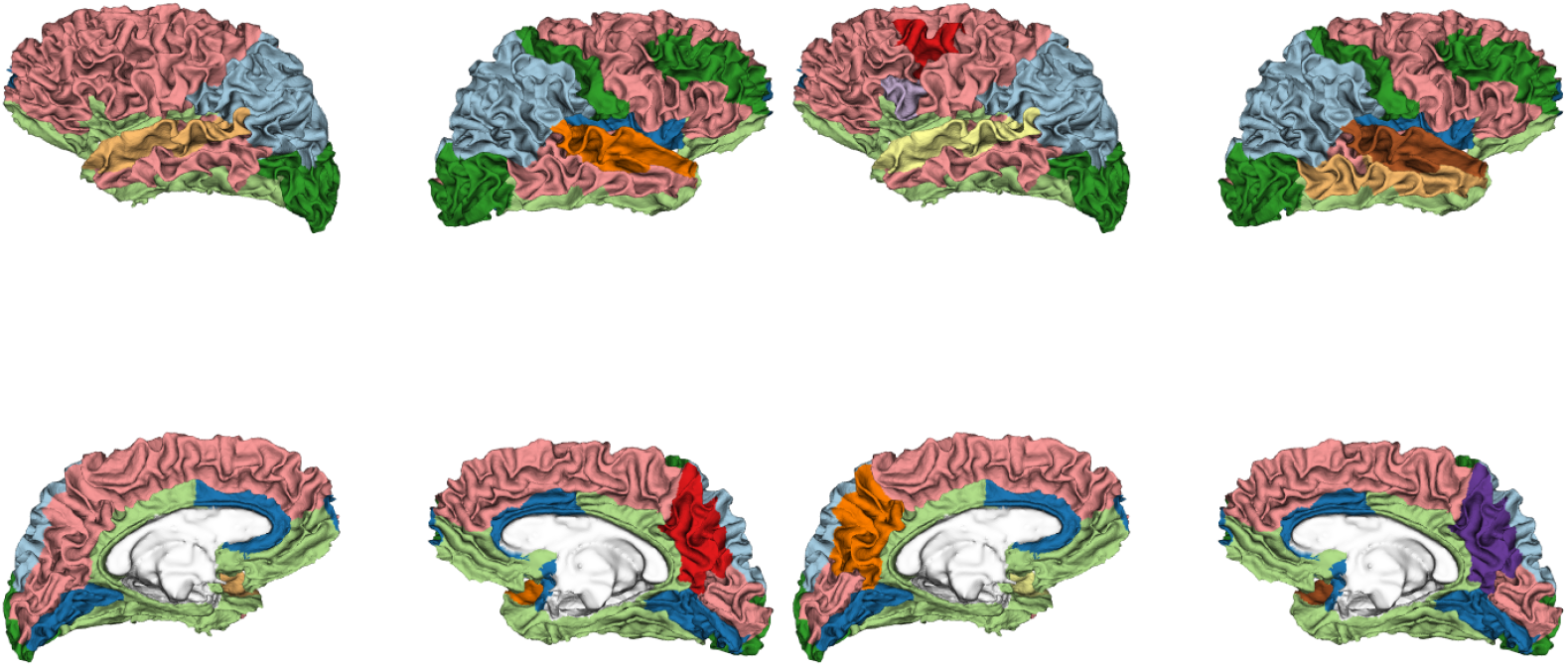
Main brain clusters on association matrix ***S***′ generated from N + → E DiffPool model predicting binary sex on unthresholded matrices. Each colour corresponds to one cluster. Left: Eight main clusters. Right: Twelve main clusters.

When looking at how the GNNs clustered the brain regions to optimise and achieve best sex prediction, we found that a clustering into 4 sets of brain regions showed interesting properties in terms of neurobiological interpretability. More specifically, the brain regions were grouped in a manner that mirrors to a certain degree the well-known cytoarchitectural and functional properties of the cerebral cortex. For example, in Figure 7 cluster 1 (dark green) included high order associative brain areas such the prefrontal and temporal cortices that have been consistently involved in complex cognitive functions such as language, working memory, and decision-making. In cluster 2, represented by the light blue colour, the GNN grouped together the left and right posterior parietal cortices which have a well-known role in visuo-spatial processing and navigation skills, amongst many other cognitive functions. Cluster 3 (in dark blue) included midline cortical areas which are part of the classic limbic or emotional system. The final cluster 4, in light green, was mainly localised in the right hemisphere and in part in the left visual cortex; it grouped together sensory-motor cortices and the right dorsolateral prefrontal cortex.

We do not wish to overinterpret our results or make “reverse neuroscience” inferences in the sense of interpreting *post hoc* the behavioural meaning of a set of regions without having directly analysed their behavioural relevance. However, we speculatively note that the clusters that discriminated binary sex the most may have some neurobiological relevance in terms of explaining the well-known behavioural differences between males and females in terms of cognitive, motor and emotional skills [45, 46]. Future work, particularly directly at investigating the links between brain and behavioural measures, is warranted to confirm whether the clustering of regions that our model has performed to achieve optimal sex classification is fully relevant to mechanistically describe the sex differences that are seen at the behavioural level.

These results demonstrate the explainability capacity of our model when using the DiffPool aggregator, which is able to cluster brain regions in a specific way for the classification task at hand.

## 5. Conclusion

In this paper we presented a novel deep learning architecture which can successfully use the high-dimensional and noisy rs-fMRI data, by leveraging not only their temporal dynamics, but also their spatial associations represented by what is commonly called the connectivity between brain locations. In contrast to previous work, we use TCNs to model temporal intra-relations and combine them with GNNs to model inter-regional relations. We illustrated and analysed the effectiveness of our model in a proof-of-concept binary sex prediction task which also included an ablation analysis with variations of the spatial pooling mechanisms. This work is, to the best of our knowledge, the first to leverage both the spatial and temporal information in rs-fMRI data in a single, end-to-end framework which includes temporal convolutions and graph neural networks, while also providing the flexibility to extract human readable explainability. Importantly, we included edge features (i.e., weights) when leveraging the graph structure in the network. This information is often ignored in the few papers which currently apply GNNs to the study of fMRI data [47]. Our ablation study showed how the graph network block was successful in leveraging the weights of the spatial dynamics, indicating the importance of designing an architecture specifically targeted for spatio-temporal rs-fMRI data. We also showed the explainability capacities of our models by analysing the clusters created by the graph hierarchical pooling mechanism, and the non-linear patterns learnt from the rs-fMRI time series.

We hope this paper can lay future groundwork on exploring flexible architectures which are able to leverage the entirety of neuromonitoring data that arise from the extremely complex spatio-temporal interplay of groups of firing neurons. Our architecture can very easily include other types of data for future work (e.g., multimodal structural and temporal data), and be extended to include possible confounds that could drive the prediction task in other brain disorders. Another exciting recent trend that can be included in our architecture is to allow the network to learn the underlying connectivity from scratch [48, 49] instead of computing associations or other handcrafted features like the ones used in this and other works [50, 51].

## Data and Code Availability

The code used to process the data from the UK Biobank is publicly available at https://github.com/ucam-department-of-psychiatry/UKB.

The code used to conduct the analysis described in this paper is publicly available at https://github.com/tjiagoM/spatio-temporal-brain.

## CRediT Authorship Contribution Statement

**T.A**.: Conceptualization, Methodology, Software, Validation, Investigation, Writing - Original Draft, Writing - Review & Editing, Visualization. **A.C**.: Conceptualization, Methodology, Writing - Review & Editing. **R.R.G**.: Resources, Data Curation. **L.P**.: Validation, Writing - Review & Editing **R.A.I.B**.: Resources, Data Curation, Writing - Review & Editing, Visualization. **P.L**.: Conceptualization, Resources, Writing - Review & Editing, Supervision, Funding acquisition. **N.T**.: Conceptualization, Methodology, Resources, Writing Review & Editing, Supervision, Funding acquisition.

## Acknowledgements

T.A. is funded by the W. D. Armstrong Trust Fund, University of Cambridge, UK. R.A.I.B. is funded by a British Academy Post-Doctoral fellow-ship and the Autism Research Trust. R.R.G is funded by the Guarantors of Brain. L.P. is funded by the Medical Research Council (MRC) grant (MR/P01271X/1) at the University of Cambridge, UK.

The UK Biobank data (application 20904) were curated and analysed using a computational facility funded by an MRC research infrastructure award (MR/M009041/1) and supported by the NIHR Cambridge Biomedical Research Centre and a Marmaduke Shield Award to Dr. Richard A.I. Bethlehem and Varun Warrier. The views expressed are those of the authors and not necessarily those of the NHS, the NIHR or the Department of Health and Social Care.

The models in this work were developed and evaluated on two different servers. Runs were performed using resources provided by the Cam-bridge Service for Data Driven Discovery (CSD3) operated by the University of Cambridge Research Computing Service (www.csd3.cam.ac.uk), provided by Dell EMC and Intel using Tier-2 funding from the Engineering and Physical Sciences Research Council (capital grant EP/P020259/1), and DiRAC funding from the Science and Technology Facilities Council (www.dirac.ac.uk). We also used Titan V GPUs generously donated to N.T. by NVIDIA.

## Footnotes

1 https://biobank.ctsu.ox.ac.uk/crystal/crystal/docs/brain_mri.pdf

2 http://surfer.nmr.mgh.harvard.edu/

3 https://nilearn.github.io/

